# The hippocampus as a perceptual map: neuronal and behavioral discrimination during memory encoding

**DOI:** 10.1101/868794

**Authors:** Manuela Allegra, Lorenzo Posani, Christoph Schmidt-Hieber

## Abstract

The hippocampus is thought to encode similar events as distinct memory representations that are used for behavioral decisions. Where and how this “pattern separation” function is accomplished in the hippocampal circuit, and how it relates to behavior, is still unclear. Here we perform in vivo 2-photon Ca^2+^ imaging from hippocampal subregions of head-fixed mice performing a virtual-reality spatial discrimination task. We find that population activity in the input region of the hippocampus, the dentate gyrus, robustly discriminates small changes in environments, whereas spatial discrimination in CA1 reflects the behavioral performance of the animals and depends on the degree of differences between environments. Our results demonstrate that the dentate gyrus amplifies small differences in its inputs, while downstream hippocampal circuits will act as the final arbiter on this decorrelated information, thereby producing a “perceptual map” that will guide behaviour.

---

To distinctly represent similar objects and events that have different behavioral significance, several brain regions are charged with the fundamental task of transforming similar inputs into well-separated neuronal representations, a process often referred to as “pattern separation” (1–4). Conversely, to recognize the same object under different contextual conditions, these brain regions need to retrieve a complete representation from degraded or incomplete inputs (5–7). This “pattern completion” process is required to organize information into categories that are equivalent in terms of their behavioral value (8).

In the context of episodic memory formation, theoretical work has suggested that these two conflicting tasks are performed by separate processes and circuits in the hippocampus (9, 10). Feedforward neurons in the input region of the hippocampus, the dentate gyrus, have been suggested to orthogonalize cortical inputs through sparse firing activity and cellular expansion (11–15). One synapse downstream of the dentate gyrus, recurrently connected pyramidal cells in CA3 are thought to retrieve memorized patterns from incomplete or degraded input via attractor dynamics (16–18). Finally, CA1, the “output” region of the hippocampus, is thought to act as a cortical translator that transfers the memory representation to the neocortex (19–21), where it is then processed to drive behavior. However, there are conflicting results about how the hippocampus encodes distinct memories of similar episodes and events, and it is unclear how these representations are used for behavioral decisions (12–15, 22–29).

To precisely control the degree of differences between spatial environments while accurately monitoring neuronal activity and behavior, we performed *in vivo* 2-photon Ca^2+^ imaging from hippocampal subregions of head-fixed mice executing a spatial memory discrimination task in a virtual reality environment (30) (Fig. 1 and Extended Data Fig. 1). Mice were initially trained to stop in a reward zone at the end of a linear virtual corridor that contained several distinct geometric objects and that was enclosed by lateral walls covered with vertical grating patterns (Fig. 1a). After successful training (~7 sessions; Fig. 1b,c: reward rate, session 1, 0.23 ± 0.5 hits per lap; session 7, 0.67 ± 0.06 hits per lap; P<0.001), mice were introduced to a novel environment with oblique grating patterns on the lateral walls but otherwise identical visual cues. To quantify the mouse’s behavioral discrimination between the memorized familiar environment (F) and the novel one (N), we moved the reward zone to the middle of the novel corridor. We initially focused our recordings on the dentate gyrus, the input region of the hippocampus (Fig. 1d–e and Extended Data Fig. 1a). We compared the firing properties of spatially modulated granule cells in the two similar but distinct environments (Fig. 1f–h). Spatially modulated place cells (20.5% (F) and 20.9% (N) of all active cells in the respective environment) differed significantly in their spatial firing patterns between the two environments (Fig. 1g: place-field correlations between odd and even lap crossings in F, 0.62 ± 0.02, n=408 cells; between F and N, 0.34 ± 0.02, n=468 cells; Fig. 1h: place-field correlations between odd and even lap crossings in F, 0.62 ± 0.02; between F and N, 0.32 ± 0.02, n=48 sessions; P<10–9). Thus, spatial population activity patterns in the dentate gyrus remapped substantially between visually similar environments.

**Figure 1.**
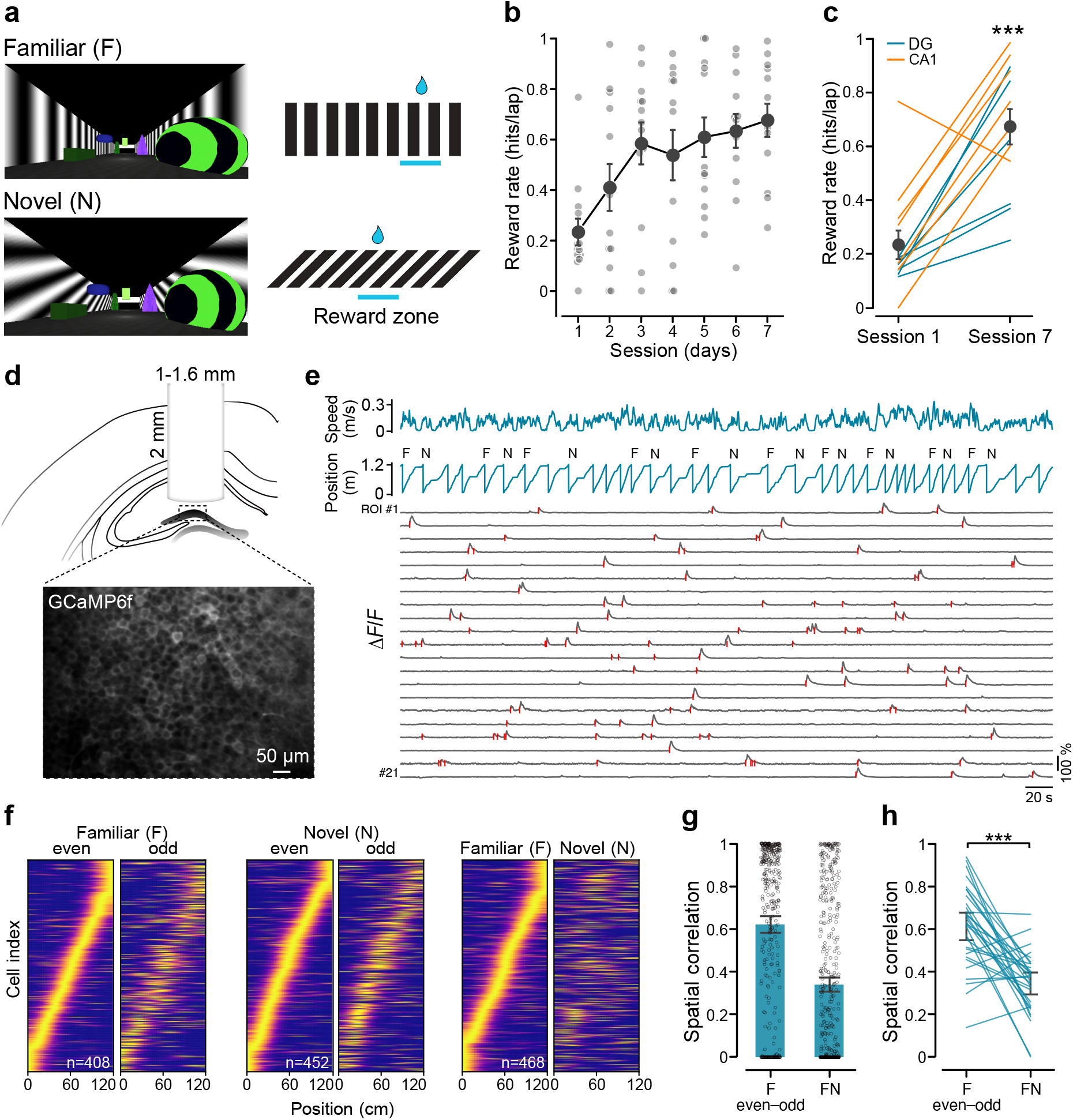
Spatial population activity in the dentate gyrus is stable and sensitive to small differences in the environment. **a,** Left, views of the familiar (vertical grating, F) and novel (oblique grating, N) environments, used for head-fixed mice navigating in virtual reality. Right, schematic indicating the corresponding reward locations along the track. **b,** Quantification of the behavioral performance, measured as reward rate (hits per lap), during the training sessions in the familiar (vertical grating) virtual environment. Light grey dots represent a single animal performance for each session, while the dark grey line and dots depict the mean ± SEM across all animals (n=13 mice). **c,** Comparison of behavioral performance during session 1 and session 7 for DG (blue lines, n=6) and CA1 (orange lines, n=7) implanted mice (day 1 vs day 7, paired t-test, P<0.001). **d,** Top, schematic drawing of the imaging window implant in the dentate gyrus. Bottom, representative fluorescence image of GCaMP6f-expressing dentate gyrus neurons *in vivo*. **e,** Representative imaging session showing the speed of the animal (blue top trace), its position along the track against time (blue bottom trace) and the context type for each lap (familiar vertical gratings or novel oblique gratings). At the bottom, fluorescence traces (in grey) extracted from example regions of interest (ROIs) for dentate gyrus neurons with significant transients indicated by red ticks (detection restricted to running periods). **f,** Spatial activity maps of all spatially modulated dentate gyrus neurons in familiar (F) or novel (N) environments sorted by the position of maximal activity (normalized per cell) in familiar even laps (left), novel even laps (middle) or all familiar laps (right) (48 sessions from 6 mice). **g,** Correlations between mean spatial activity maps across spatially modulated cells in the same familiar (F even–odd, left bar) and in different environments (FN, right bar) in the dentate gyrus (F even–odd, n=408 cells; FN, n=468 cells). **h,** Correlations between mean spatial activity maps across recording sessions in the same familiar (F even–odd, left bar) and in different environments (FN, right bar) in the dentate gyrus (n=48 sessions recorded from 6 mice; t-test across sessions, P<10^−9^). *** P<0.001.

We next sought to examine how this orthogonalised information is processed downstream in the output region of the hippocampus, CA1 (Fig. 2 and Extended Data Fig. 1b). Despite the pronounced spatial remapping in the dentate gyrus, spatial correlations between different environments were substantially larger in the downstream region CA1 (Fig. 2d: place-field correlations between odd and even lap crossings of the familiar environment, 0.56 ± 0.01, n=1133 cells; between familiar and novel environments, 0.46 ± 0.01, n=1406 cells; Fig. 2e: place-field correlations between odd and even lap crossings of the familiar environment, 0.54 ± 0.02, between familiar and novel environments, 0.47 ± 0.02, n=29 sessions; P<0.05). As a consequence, spatial *de*-correlation (i.e. the reduction in correlation; see Methods) between familiar and novel environments was significantly more pronounced in the dentate gyrus than in CA1 (decorrelation in DG: 0.31 ± 0.04, n=48 sessions; CA1: 0.07 ± 0.03, n=29 sessions; P<10^−4^; Fig. 2f). Thus, our data suggest that small visual differences between environments are sufficient to result in pronounced pattern separation in the dentate gyrus, but fail to induce remapping in the downstream circuit CA1, where pattern completion leads to similar spatial representations of the familiar and novel environments.

**Figure 2.**
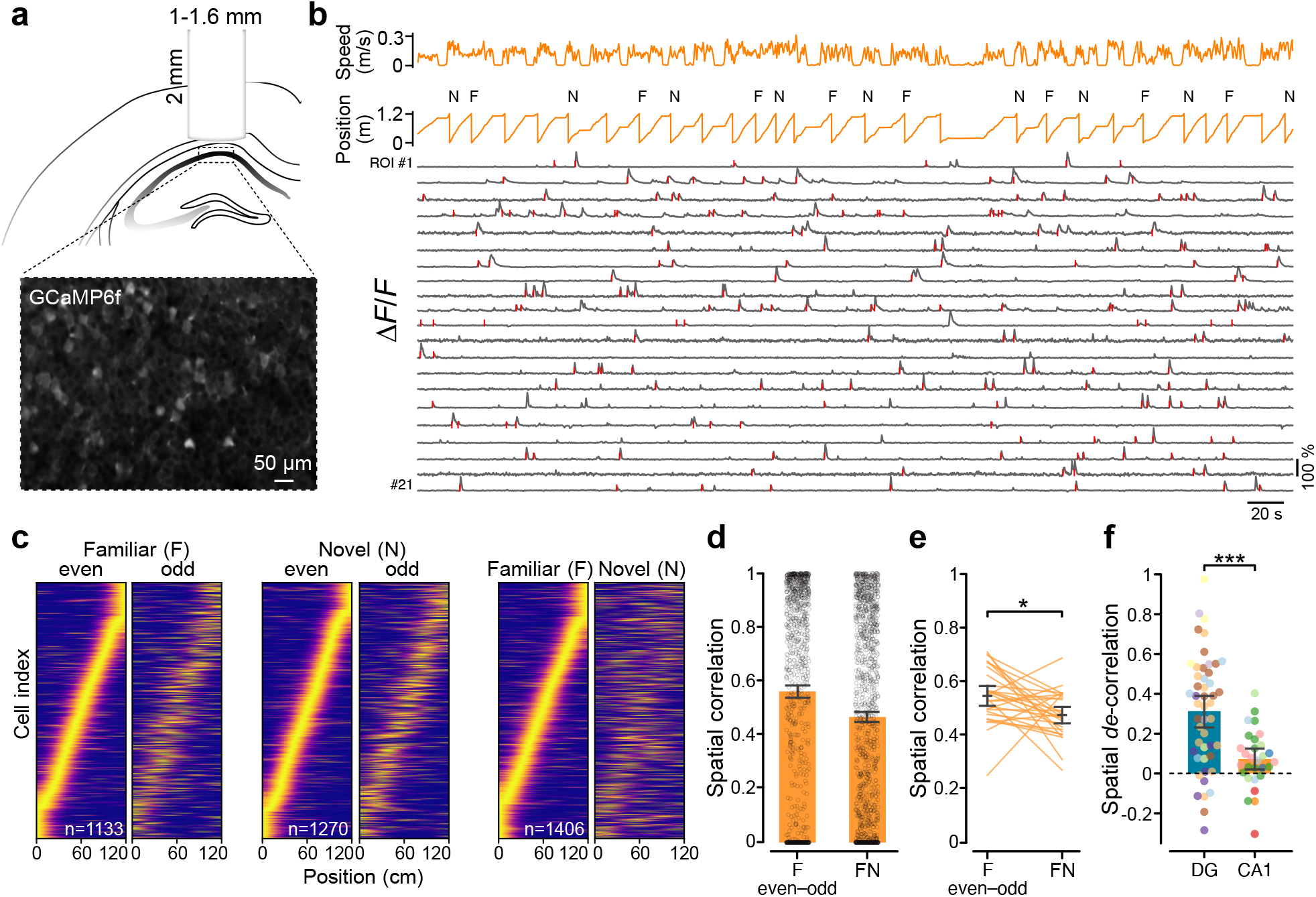
Small differences in environments lead to less spatial remapping in CA1 than in the dentate gyrus. **a,** Top, schematic drawing of the imaging window implant in CA1. Bottom, representative fluorescence image of GCaMP6f-expressing CA1 neurons *in vivo*. **b,** Example of a representative imaging session showing animal speed and position (orange lines), and fluorescence traces (grey) extracted from a subpopulation of CA1 pyramidal neurons, with significant calcium transients indicated by red ticks (detection restricted to running periods). **c,** Spatial activity maps of all spatially modulated CA1 neurons in familiar (F) or novel (N) environments sorted by the position of maximal activity in familiar even laps (left), novel even laps (middle) or all familiar laps (right) (29 sessions from 7 mice). **d,** Correlations between mean spatial activity maps across spatially modulated CA1 neurons in the same familiar (F even–odd, left bar) and in different environments (FN, right bar) (F even– odd, n=1133 cells; FN, n=1406 cells). **e,** Correlations between mean spatial activity maps across recording sessions in the same familiar (F even–odd, left bar) and in different environments (FN, right bar) in CA1 (n=29 sessions recorded from 7 mice; t-test across sessions, P<0.05). **f,** Spatial *de*-correlation, quantified as the difference between spatial correlations within the same familiar (F even–odd) and different environments (FN), in the dentate gyrus (DG) and in CA1. Dots indicate single recorded sessions from different animals (DG, n=48 sessions from 6 mice; CA1, n=29 sessions from 7 mice; t-test, P<10^−4^). * P<0.05; *** P<10^−4^.

In addition to physical space, the hippocampus also encodes non-spatial dimensions and experiences (31, 32). To assess neuronal discrimination beyond spatial remapping, we sought to analyse neuronal discrimination of the different contexts (vertical vs. oblique gratings) independently of the spatial position of the animal (Fig. 3). We first quantified how selectively neurons, including all place and non-place cells, fired in the familiar or the novel environment (Fig. 3a,b). Our analysis revealed that dentate gyrus neurons were significantly more selective than CA1 neurons (selectivity in dentate gyrus: 0.72 ± 0.01, n=3289 active cells; CA1: 0.59 ± 0.00, n=10587 active cells; P<10–69; Fig. 3b). The higher selectivity in dentate gyrus was not only a consequence of its overall lower activity levels, as selectivity in the dentate gyrus was higher than in CA1 throughout the whole range of observed activity rates (Fig. 3c). To quantify more rigorously how informative population activity is about the environment, we established a binary decoder that uses activity rates to classify population vectors into one of the two environments (33). We found that the decoder was significantly more successful at classifying dentate gyrus than CA1 activity (F-N decoder performance (AUC) in the dentate gyrus: 0.67 ± 0.02, n=59 sessions; CA1: 0.55 ± 0.01, n=32 sessions; P<0.001; chance level = 0.5; Fig. 3d). Thus, independent of positional information, activity in the two similar environments was more orthogonalised in the dentate gyrus than in CA1. These results are consistent with the notion that, in addition to spatial remapping, the dentate gyrus uses sparse selective coding to achieve pattern separation (13).

**Figure 3.**
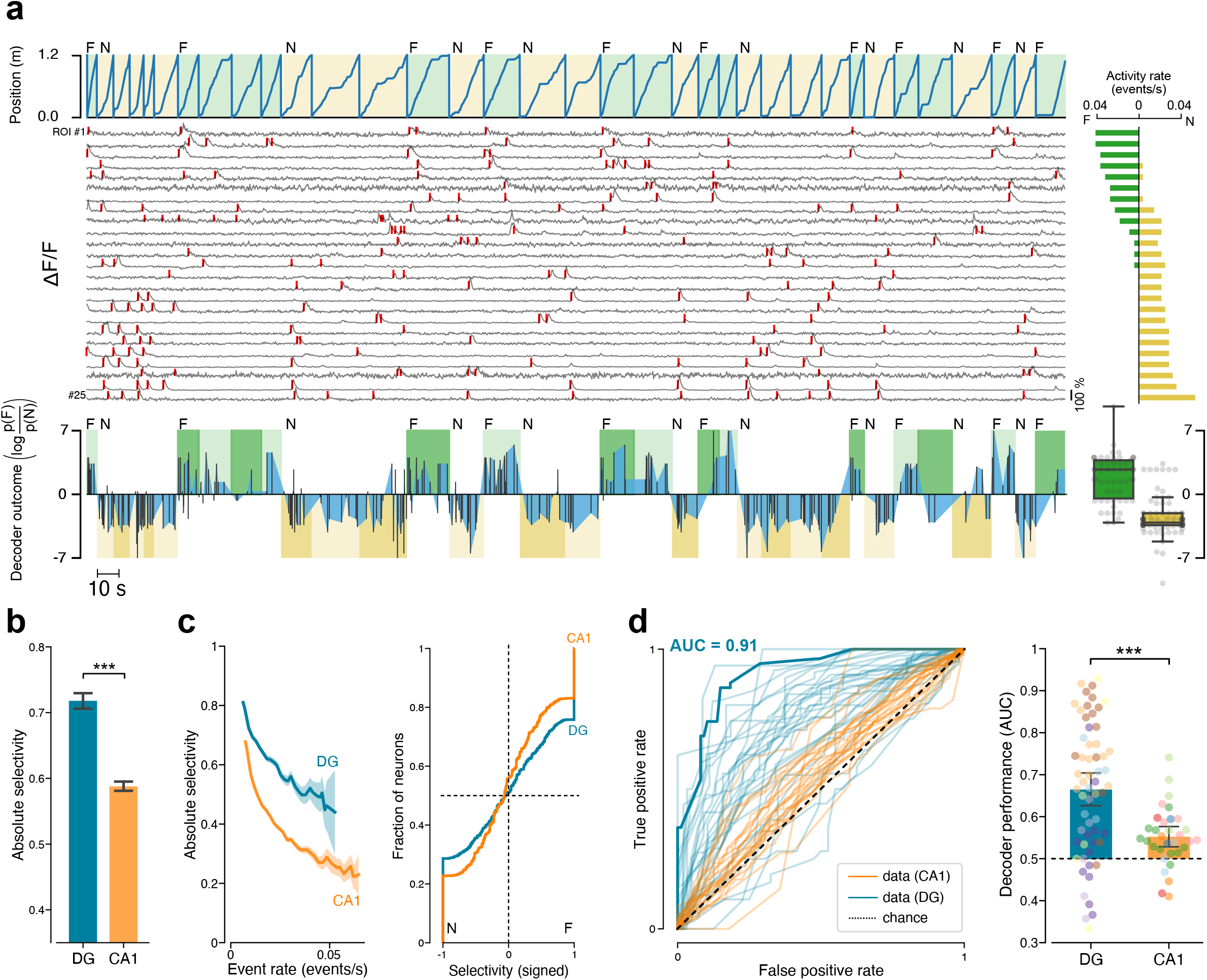
The dentate gyrus discriminates between small changes in contextual information. **a,** Top: Example session recorded from the dentate gyrus (DG) showing the position of the animal against time, the context type for each lap (F, familiar vertical gratings in green; N, novel oblique gratings in yellow) and fluorescence traces extracted from the 25 most active neurons (grey), with significant transients indicated by red ticks (detection restricted to running periods). For each fluorescence trace on the left, its event rate in the familiar (green) and novel (yellow) environments is reported on the right. Bottom: decoder outcome for the session shown at the top. Light and dark color shades refer to training and test data, respectively (same color code as in top panel). Black bars connected by blue areas represent the decoder outcome (difference in log-likelihood of the two environments; F: positive; N: negative). Bottom right: Summary of decoder outcome for the test data of the example session. Data points represent time bins. **b,** Absolute selectivity (see Methods) for the two environments F and N in DG (blue) and CA1 (orange) cells (DG: n=3289 active cells; CA1: n=10587 active cells; P<10^−69^). **c,** Left: Absolute selectivity for DG and CA1 neurons plotted against their firing rate. Right: Cumulative histogram where each neuron was scored depending on its selectivity towards the familiar F (positive) or the novel N (negative) environment. **d,** Left: Quantification of the decoder performance (Receiver operating characteristic, ROC). Blue traces: DG sessions; orange traces: CA1 sessions. The dark blue line refers to the recording session shown at the top of panel **a**. Right: Comparison of decoder performance, quantified as area under the ROC curve (AUC), between the dentate gyrus and CA1. Dots represent single recorded sessions from different animals (DG, 59 sessions from 6 mice; CA1, 32 sessions from 7 mice; t-test, P<0.001; chance level 0.5). *** P<0.001.

Previous work has shown that spatial coding in CA1 may be affected by task relevance and engagement (34–38). As neuronal pattern separation is thought to underlie behavioral discrimination, we examined how spatial and non-spatial decorrelation in the dentate gyrus and CA1 relate to behavioural discrimination (Fig. 4). Addressing this question required us to explore a large range of degrees of neuronal discrimination in both the dentate gyrus and in CA1. Therefore, we introduced another novel environment (N*) that differed substantially from the familiar one (Fig. 4a; Extended Data Fig. 2–6). To assess behavioural discrimination, the reward zone was placed at the center of N*. Animals showed higher performance in discriminating between F and N* than between F and N (Fig. 4c: reward rate in N, 0.26 ± 0.03; in N*, 0.37 ± 0.04, n=37 sessions, P<0.01; Extended Data Fig. 6a). This increase in behavioral performance was accompanied by a larger neuronal discrimination in CA1, but not in the dentate gyrus (Extended Data Fig. 4). These data indicate that neuronal discrimination in CA1, but not in the dentate gyrus, reflects behavioral discrimination between the two environments. Further supporting this notion, we found that in CA1, but not in the dentate gyrus, all measures of behavioral and neuronal discrimination were positively correlated (Fig. 4d: DG, n=13, P>0.05; CA1, n=41, P<0.01; Extended Data Fig. 6). Consistently, behavioral discrimination negatively correlated with the neuronal “confusion” (i.e. the spatial correlation between the two different environments) in CA1, but not in the dentate gyrus (Fig. 4e: DG, n=13, P>0.05; CA1, n=41, P<0.01). Together, our data suggest that neuronal discrimination in the dentate gyrus can occur below the animal’s threshold to perceive behaviorally relevant changes in the environment, while representations in CA1 reflect the behavioral decision of the animal.

**Figure 4.**
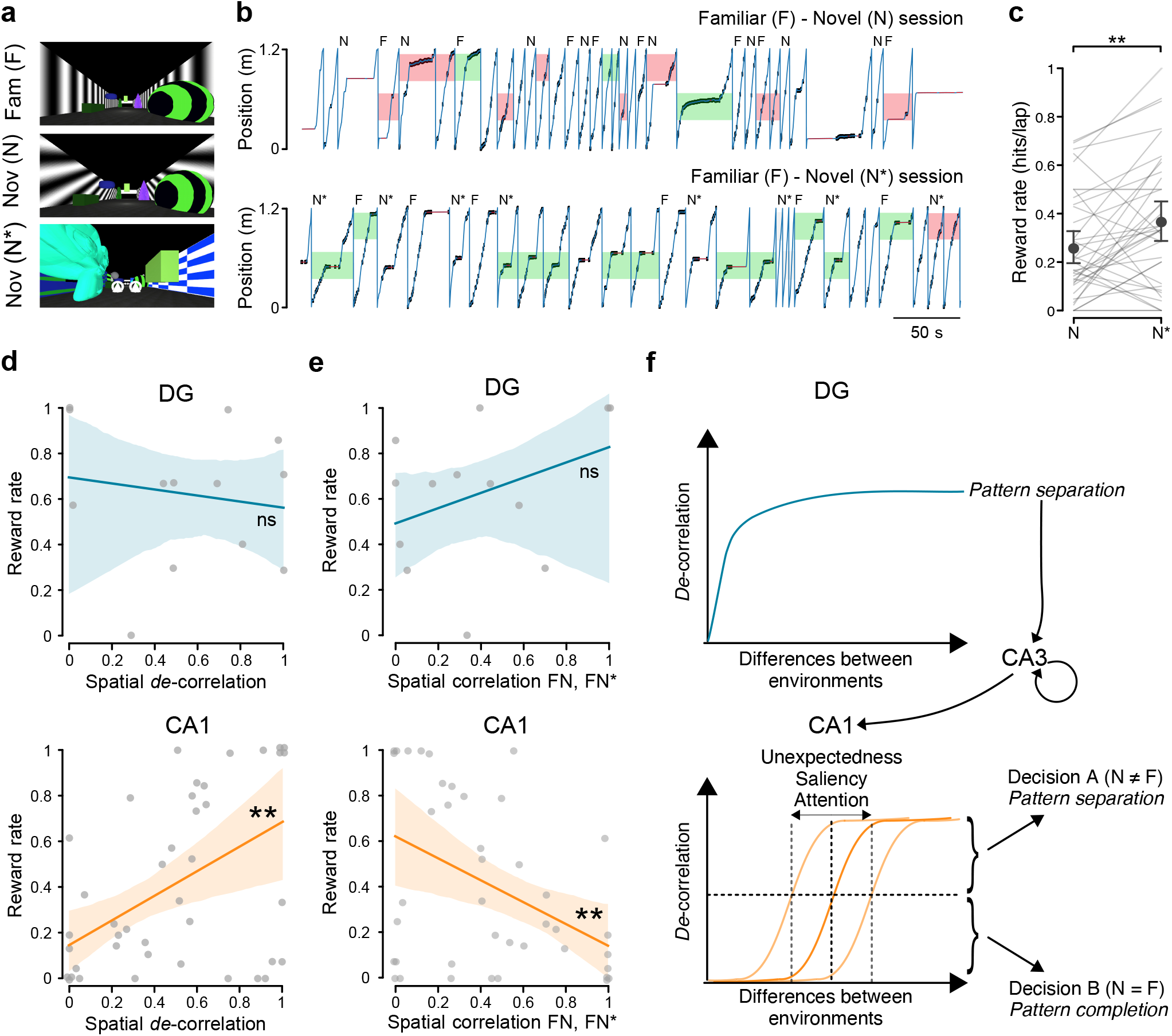
Neuronal discrimination in CA1, but not in the dentate gyrus, reflects behavioral discrimination. **a,** Views of the virtual environments (Fam (F), familiar; Nov (N) or (N*), novel). **b,** Illustration of two representative recording sessions in the familiar F - novel N (top) and familiar F - novel N* (bottom) environments. The blue traces represent the position of the animal along the track and the black ticks indicate animal licks. The green rectangles indicate the correct licking choice in the familiar (F) and novel (N or N*) laps, while the red blocks refer to the false choice, i.e. the animal licking in the wrong reward zone, for the familiar (F) and novel (N or N*) laps. **c,** Quantification of behavioral performance in the novel (N and N*) environments, measured as reward rate (hits per lap; n=37 sessions; P<0.01). **d,** Pearson’s correlation between behavioral performance (reward rate) and spatial *de*correlation in DG (top; 13 sessions from 2 mice; P>0.05) and CA1 (bottom; 41 sessions from 7 mice; P<0.01). Dots indicate single recorded sessions from different animals. **e,** Pearson’s correlation between behavioral performance (reward rate) and spatial correlation between familiar (F) and novel (N or N*) environments in DG (top; 13 sessions from 2 mice; P>0.05) and in CA1 (bottom; 41 sessions from 7 mice; P<0.01). **, P<0.01. **f**, Schematic: The dentate gyrus generates robustly decorrelated neuronal representations of different environments, even if the differences are very small (top). This decorrelated information is transmitted to CA1, which shows either pattern completion or pattern separation by integrating it with several additional inputs mediating the unexpectedness and saliency of the environment. The representation in CA1 is then used by downstream neocortical circuits to guide behavioral decisions. CA1 thereby produces a “perceptual map” of the environment: it remaps in response to changes in sensory inputs that exceed a saliency threshold so that the animal perceives them as behaviorally relevant.

To rule out that neuronal discrimination between different environments could be performed based on visual inputs alone, we passively presented previously recorded virtual reality sessions to the animals in an open-loop paradigm. These experiments confirmed that neuronal discrimination in both the dentate gyrus and CA1 required active engagement of the animal in a navigational task, as we found a significant decrease of neuronal discrimination in both regions when the passive movie was presented (Extended Data Fig. 7).

We conclude that the dentate gyrus produces highly orthogonalised output by activating nonoverlapping subpopulations of neurons in response to small changes between a memorized familiar and a novel environment. By contrast, neuronal discrimination in CA1 shows spatial remapping when differences between environments are larger. At the same time, the magnitude of neuronal discrimination in CA1, but not in the dentate gyrus, closely reflects the behavioral performance of the animal.

It has been suggested that hippocampal circuits downstream of the dentate gyrus perform a “pattern completion” operation by virtue of their recurrent connectivity (16), whereby representations fall into one of several “attractor” states that are robust to degraded or incomplete input information. These attractor properties may explain why population activity in CA1 sharply remaps from one representation to the other once the differences between environments are sufficiently large, while the dentate gyrus feedforward circuit produces a more constant decorrelation throughout a range of differences between environments.

Which mechanism can explain that the dentate gyrus performs constant decorrelation, while the downstream circuits appear to remap independently? We suggest that the robustly decorrelated inputs from the dentate gyrus are pushing downstream circuits towards forming a new representation, whereas their attractor dynamics counteract this process by stabilizing the current map. Additional inputs, such as information about the saliency and the unexpectedness of the novel environment, are required for CA3 and CA1 to form a new representation. Thus, the level of saliency decides whether the decorrelated information from the dentate gyrus can succeed to shift CA3 and CA1 cell assemblies towards a new attractor state (Fig. 4f).

CA1 receives several streams of inputs, such as orthogonalized information from the dentate gyrus, which is transmitted to CA1 via CA3; direct input from the entorhinal cortex via the perforant path, that is required for precise positional and contextual information; and inputs that convey the unexpectedness and the saliency of the environment, and consequently the animal’s attentional and arousal state. Future experiments will have to reveal the precise relative importance of these sources to the remapping threshold in CA1, and the neuronal origins of the saliency signal. Candidate input streams include neuromodulatory systems, such as cholinergic inputs from the medial septum indicating novelty and attention (39), or noradrenergic inputs from the Locus coeruleus mediating arousal. These different contributions to the remapping threshold in CA1 may explain why previous work on neuronal discrimination in different hippocampal subregions has provided inconsistent results (12, 13, 15, 22–27, 29), as our data suggest that the behavioral relevance of the navigational task plays a crucial role in neuronal discrimination.

In summary, our data reveal how orthogonalised neuronal memory representations in the input and output regions of the hippocampus can drive behavioral discrimination. We propose that the dentate gyrus robustly reports changes in external and internal variables, even if the animal does not perceive the differences as behaviorally relevant. By contrast, the output region CA1 produces a “perceptual map” of the environment: it remaps in response to changes in sensory inputs that exceed a saliency threshold so that the animal can perceive them as behaviorally relevant. Downstream neocortical circuits then use this perceptual map to drive behavioral decisions.

## Acknowledgments

We thank Josef Bischofberger, Rémi Monasson, Yaniv Ziv, and David DiGregorio for helpful discussions and comments on the manuscript. We thank Lucile Sontag and Claire Lecestre for their technical support. This work was supported by grants from the ERC (StG 678790 NEWRON to C.S.-H.), the Pasteur Weizmann Council, and a Pasteur-Roux fellowship to M.A.

## METHODS

### Mice

All procedures were carried out in accordance with the national guidelines on the ethical use of animals of EU Directive 2010/63/EU and were approved by the Ethics Committee CETEA of Institut Pasteur (protocol number 160066). Adult C57BL/6J male mice (Janvier Labs) were used for all experiments at an age of 6-16 weeks. After their arrival at 4-5 weeks old, mice were housed in a room with a temperature of 21°C, maintained on a 12/12 h light-dark cycle and grouped in 2-4 in cages enriched with running wheels. Food and water were available *ad libitum* until the beginning of the experiments, when they were placed under controlled water supply (0.5 mg of hydrogel per day, Clear H2O, BioService) and maintained at 80-85% of their initial body weight over the course of imaging experiments. In total, imaging data from 13 mice (DG, n=6; CA1, n=7) were used in this study.

### Surgery

All surgical procedures were performed in a stereotaxic apparatus (Kopf instruments). Mice were maintained under isoflurane anaesthesia (5% for induction and 1.5% for surgery, vol/vol) and analgesia (0.05 mg/kg buprenorphine, Vetergesic). During the surgery, mice were kept at a body temperature of ~36°C using a heating blanket, and their eyes were protected with artificial tear ointment. Postoperative analgesic treatment was continued with metacam (1 mg/kg, Méloxicam) dissolved in a recovery diet gel (BioService) for 2 days after surgery.

### Stereotaxic injections

After disinfection with Betadine, the skin was opened and the exposed cranial bone was cleaned of connective tissue. A small craniotomy was performed above the right dorsal hippocampus (2 mm posterior to bregma and 1.5 mm lateral), and 500 nL of AAV1.Syn.GCaMP6f.WPRE.SV4 (titre 3.4×10^12^ TU/mL; University of Pennsylvania Vector Core) were injected via a glass micropipette (Wiretrol, 5-000-1010 Drummond) at a depth of 1.7 mm from the dural surface for targeting the DG, and 1.2 mm for CA1. Mice were allowed to recover for at least 2 days after injection before undergoing any subsequent procedure.

### Chronic imaging window and head plate implantation

To record Ca^2+^-dependent fluorescence changes of the GCaMP6f sensor, we used two different imaging implant strategies, either a graded index lens (GRIN; 1 mm diameter, 3.4 mm height, NA = 0.5, G2P10 Thorlabs), or a custom made imaging cannula. The imaging cannula was assembled by UV curing (Norland optical adhesive) a circular coverglass (1.6 mm diameter, 0.16 mm thickness, produced by Laser Micromachining) to a cylindrical stainless steel tube (2 mm height, 1.65 mm outer and 1.39 mm inner diameter; Coopers Needleworks). Imaging window implantations were performed several days after the initial viral injection. Mice were anaesthetised as described above and, after the scalp was exposed, a craniotomy (~1 or ~1.6 mm diameter) was made centred on the previous injection site. For cannula implants, the overlying cortex (including parts of the somatosensory and posterior parietal association cortices) was gently aspirated with a 27 gauge needle while constantly being rinsed with aCSF solution (29, 40), and bleeding was controlled with a gel dental sponge (Gelfoam, Pfizer). We terminated the aspiration when the external capsule became visible. The outer part of the external capsule was then gently peeled away using a fine forceps, leaving the inner capsule and the hippocampal formation itself undamaged for CA1 imaging recordings. To provide optical access to the dorsal blade of the DG, we continued to gently aspirate CA1 directly superior to the dentate gyrus until the loose fibers and vasculature of *stratum moleculare* were visible. For GRIN lens implants, a beveled stainless steel cylinder (1mm diameter) was slowly lowered to the target region without any cortical aspiration, and removed before proceeding with the implant. The window (cannula or GRIN lens) was then gradually lowered into the craniotomy until the tip was in contact with the internal capsule for CA1 imaging, or 100-200 µm above the hippocampal fissure for dentate gyrus imaging. Protruding parts of the GRIN lens or cannula were secured to the skull with opaque dental cement (Super-bond C&B, Phymep). Neuronal activity was largely comparable between the two optical window strategies (Fig. S3). Mice were then implanted with a stainless-steel headpost for head fixation (Luigs & Neumann) during imaging experiments. Finally, the conical portion of a nitrile rubber seal (749-575, RS Components) was glued to the head-plate with dental cement and filled with a silicone elastomer (900-2822, Henry Schein) to protect the window preparation during recovery and between recording sessions.

### Virtual-reality environment

A virtual reality setup was implemented as previously described (30). Briefly, head-fixed mice navigated on a cylindrical polystyrene treadmill (20 cm in diameter), rotating forward or backward. Cylinder rotation associated with animal locomotion was read out with a computer mouse (G203, Logitech) at a poll rate of 1KHz and converted to a movement in a rectangular virtual reality scene. The virtual environments were displayed onto a spherical dome screen (240°), covering nearly the entire field of view of the animal. Below the mouse, a quarter-sphere mirror (45 cm diameter) and a projector (Casio XJ-A256) were used to project the virtual reality scenes on the dome. The virtual linear corridor was 1.2 m long, with objects placed along the linear track and vertical or oblique grating textures on the lateral walls. A reward zone was located within each linear virtual track, at the end (0.81-1.15 m) or at the centre (0.34-0.68 m) of the vertical and oblique corridors, respectively. An enriched environment, used to drive better behavioural discrimination and neuronal discrimination in CA1, consisted in a substantially different virtual scene, with different visual cues and textures on the walls, but maintaining the same reward location as for the oblique corridor. The Blender Game Engine (www.blender.org) was used in conjunction with the Blender Python API to drive the virtual reality system.

### Behavioural training and analysis

Two weeks after the imaging window implantation, water restricted mice were handled 10 min per day for 3 days and placed on the treadmill for 10-20 min for 2 consecutive days to get habituated to both the experimenter manipulation and the experimental setup. After the habituation, mice underwent 7 training sessions, 30 min each, over the course of 1-2 weeks before recordings (Fig. 1b-c). Mice were trained to run along the linear virtual corridor. A drop of sugar water (10 µl, 8 mg/mL sucrose) was dispensed by a spout placed in front of their mouth as a reward if they spent 2 s or more within the reward zone. When the animals reached the end of the linear track, they were “teleported” back to the start of the virtual corridor after crossing a black frontal wall, indicating the end of a lap and the onset of the subsequent one. No punishments were provided in our experimental design. During the training, only the familiar (vertical gratings) environment was displayed, while from the first day of imaging sessions, mice were presented with a random alternation of familiar and novel environments. In particular, each imaging session was organised into about 10 min of recording, where similar environments (familiar: vertical gratings (F) vs novel: oblique gratings (N)) or substantially different environments (familiar: vertical gratings (F) vs novel: enriched environment (N*)) were randomly displayed. Training and imaging recordings were performed during the dark cycle of the mice. Behavioral performance was comparable between animals implanted for dentate gyrus or CA1 imaging (Fig. S8).

### In vivo two photon calcium imaging

*In vivo* imaging was performed using a resonant-galvanometer high speed laser-scanning two-photon microscope (Ultima V, Bruker), with a 16x, 0.8 NA water immersion objective (Nikon). Time-series images were acquired at 30 Hz frame rate (512×512 pixels, 0.8–1.2 µm/pixel), with a femtosecond-pulsed excitation laser beam (Chameleon Ultra II, Coherent) tuned to 920 nm for imaging GCaMP6f expressing cells. To block diffused light from the projection system, a black foam rubber ring was positioned between the animal’s implant and the objective, and a green light blocking filter (FES0450, Thorlabs) was placed in front of the projector light output.

### Histology for imaging area detection

At the end of the experiments, mice were deeply anaesthetized with ketamine/xylazine and transcardially perfused with PBS followed by 4% paraformaldehyde/PBS. Brains were removed and post-fixed overnight in 4% paraformaldehyde/PBS. Brains were then cut into 60 µm coronal slices, sections containing the area underneath the imaging window were collected, and the correct positioning of the imaging cannula or GRIN lens was confirmed by fluorescence microscopy.

### Data analysis

#### Imaging data processing

Motion correction of the imaging data was performed using a phase-correlation algorithm built into the suite2p software (41) (http://github.com/cortex-lab/Suite2P). Segmentation into regions of interest (ROIs) was performed with suite2p using a singular value decomposition algorithm. To select dentate gyrus granule cells, we only used ROIs corresponding to small, densely packed cell bodies. ROIs corresponding to large isolated cell bodies were discarded in order to avoid interneurons and mossy cells. The neuropil signal was subtracted from the extracted fluorescence using suite2p. Neuronal activity was quantified by adapting previously published methods (40): “Events” were identified as contiguous regions in the d*F*/*F* signal exceeding a threshold of mean +2.5 standard deviations of the overall d*F*/*F* signal, and exceeding an integral above threshold of 7,000 d*F*/*F*.

#### Place cell identification

To compute spatial activity maps, we used data from continuous running periods with a duration > 1 s at a speed > 0.5 cm/s. For each lap crossing, spatial maps were computed by dividing the track into 50 spatial bins and then smoothing the occupancy vector of events with a gaussian kernel (sigma = 5 cm).

We defined spatially-modulated cells as those neurons that fire consistently at the same location between different lap crossings, independently of the exact shape of their place fields. Therefore, spatially modulated cells were selected based on the mean cross-correlation between their spatial maps in different lap crossings. For each neuron, we dissociated firing activity and spatial position by shuffling the recorded position in chunks of 300 ms, and repeated this bootstrap procedure 100 times to obtain a null model for the mean cross correlation. Those neurons whose mean cross-correlation exceeded the bootstrap with a Z-score higher than 2.0 were identified as place cells.

#### Rate vector and selectivity

The event rate for the i-th cell was defined as the number of identified neural events of a cell divided by the total running time in a lap or in a session. For each session, selectivity of the i-th neuron was defined as the normalized difference of event rate of that neuron computed during familiar and novel lap crossings (referred to as A and B, respectively): 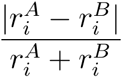. “Signed” selectivity was defined accordingly, with no absolute value at the numerator. The rate vector was defined as the collection of event rates of a population of neurons during a lap crossing or a session.

#### Neural discrimination between contexts

Neural discrimination between contexts (referred to as “decorrelation”) was computed by comparing the similarity of population activity during lap crossings of the familiar environment (F even-odd) with the similarity of population activity computed between different environments (familiar-novel, FN). To compute F even-odd correlations, we divided the exploration of familiar environment into odd and even lap crossings, and computed the cross correlation of place fields of neurons in the two cases. We used cells active in the “even” crossings for the comparison. To compute FN correlations, we compared place fields computed during exploration of the familiar and novel lap crossings. Cells active in the familiar environment were used for the comparison.

#### Inference-based decoder for environmental representation

To decode the environmental representation from neural activity alone (i.e., with no information on the precise location of the animal on the track), we employed published methods based on probabilistic modeling of population activity and Bayesian hypothesis testing (33). In brief, two statistical models, one for each environmental condition, were inferred from two collections of population vectors (discretized and binarized events in a time window of 120 ms) recorded during exploration of the two corresponding environments, here denoted as *A* and *B*. By defining the binarized activity of the i-th cell in a time bin as *s*_*i*_ ∈ 0, 1, each model is a probability distribution that scores any activity vector s = (*s*_1_, *s*_2_, …, *s*_*N*_) given the environmental variable *M* ∈ {*A, B*}. The model reads

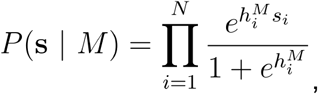

where parameters 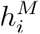 control the mean event rate of the i-th neuron and are inferred such that the probabilistic model reproduces, on average, the mean event rate observed in the training set. For this particular class of models (statistically-independent neurons), this procedure can be carried out analytically and yields

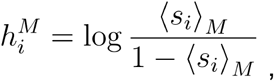

where the notation 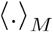 indicates the average over the population vectors in the training set for environment *M*.

We then decoded the environmental variable *M* from the *test* population vectors by comparing the two probabilistic models in a Bayesian hypothesis test. For each test vector s, we computed the log-likelihood difference between the two environments as

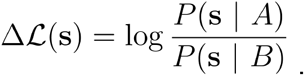

A positive value of 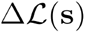 indicates that the test activity vector is more likely to have been sampled from environmental conditions *A* than *B*, and vice-versa for negative values.

#### Non-spatial decorrelation: performance of the environment decoder

Events were discretized into binary population activity vectors corresponding to time windows of 120 ms. Lap crossings in a recording session were then divided according to the two environmental conditions, and then further into training and test data. Activity vectors corresponding to even lap crossings were used to train the binary decoder, which was used to predict the environmental conditions of activity vectors sampled during *odd* lap crossings. For each test vector, we computed the decoder outcome 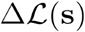 and assessed the performance of the decoder on the population of 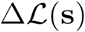 values on a session basis. The performance of the decoding procedure was assessed by contextualizing the 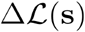 signal within the binary decoder theory and receiving-operator characteristic curves (42), as established in previous publications (33, 43). In brief, a threshold 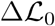 is chosen, and population vectors whose delta log-likelihood difference 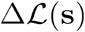 exceeded the threshold were classified as “positive” (corresponding to environment A), while the ones below were classified as “negative” (corresponding to environment B). The threshold value is varied in a large interval, and for every value we compute the fraction of correctly-classified A patterns (true positive rate, TPR) and the fraction of falsely-classified A patterns (false positive rate, FPR). Each pair of TPR-FPR draws a point on the so-called ROC curve. The area under the ROC curve (AUC) is taken as a measure of performance.

### Behavioral discrimination between contexts

For each session, we used the lick behaviour of the animal in the novel environment to quantify behavioral discrimination. If the animal confuses the novel environment for the familiar one, we expect it to lick in anticipation for a reward in the reward zone corresponding to the wrong (familiar) environment, i.e., at the end of the track. Likewise, we expect the animal to lick only in the correct reward zone (novel, center of the track) when it recognizes the novel environment and learns the task. To quantify behavioral discrimination, for each lap crossing we detected a *hit* if the ratio between number of licks in the correct (novel) reward zone and the region of the track outside any reward zone was higher than a threshold (set to 1.2). Likewise, an *error* was recognized if the ratio between the number of licks in the wrong (familiar) reward zone and the region of the track outside reward zone was higher than the same threshold. d-prime was computed for each session by using all errors and hits of the corresponding lap crossings, excluding those laps where both an error and a hit were recognized.

### Statistics and visualization

All values are given as mean ± SEM, unless stated otherwise. Statistical significance was assessed using either paired or unpaired Student’s t tests, as appropriate. In some figures, a small amount of jitter was applied to coinciding data points to improve visual clarity.

