## Supplemental Material for "The hippocampus as a perceptual map: neuronal and behavioral discrimination during memory encoding"

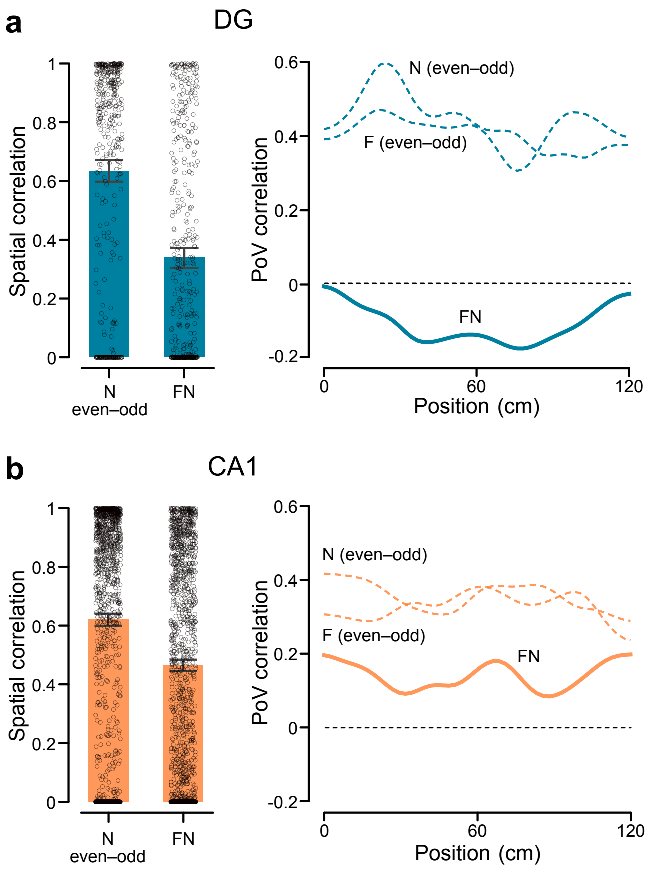


**Extended Data Figure 1**: Spatial and population vector activity correlations in the same (novel N) or different (familiar F - novel N) environments in the dentate gyrus and CA1

**a,** Supplement to Fig. 1f. Left: Correlations between mean spatial activity maps across spatially modulated cells in the same novel (N even–odd, left bar) and in different environments (FN, right bar) in the dentate gyrus (N even–odd, 0.63 ± 0.02, n=452 cells; FN, 0.34 ± 0.02, n=468 cells; 48 sessions recorded from 6 mice). Right: Population vector correlation (PoV) across mean spatial activity maps for all spatially modulated cells against the position of the animals in the virtual corridor. **b,** Supplement to Fig. 2c. Same analysis as in **a**, but for CA1 (left; N even–odd, 0.62 ± 0.01, n=1270 cells; FN, 0.47 ± 0.01, n=1406 cells; 29 sessions recorded from 7 mice).

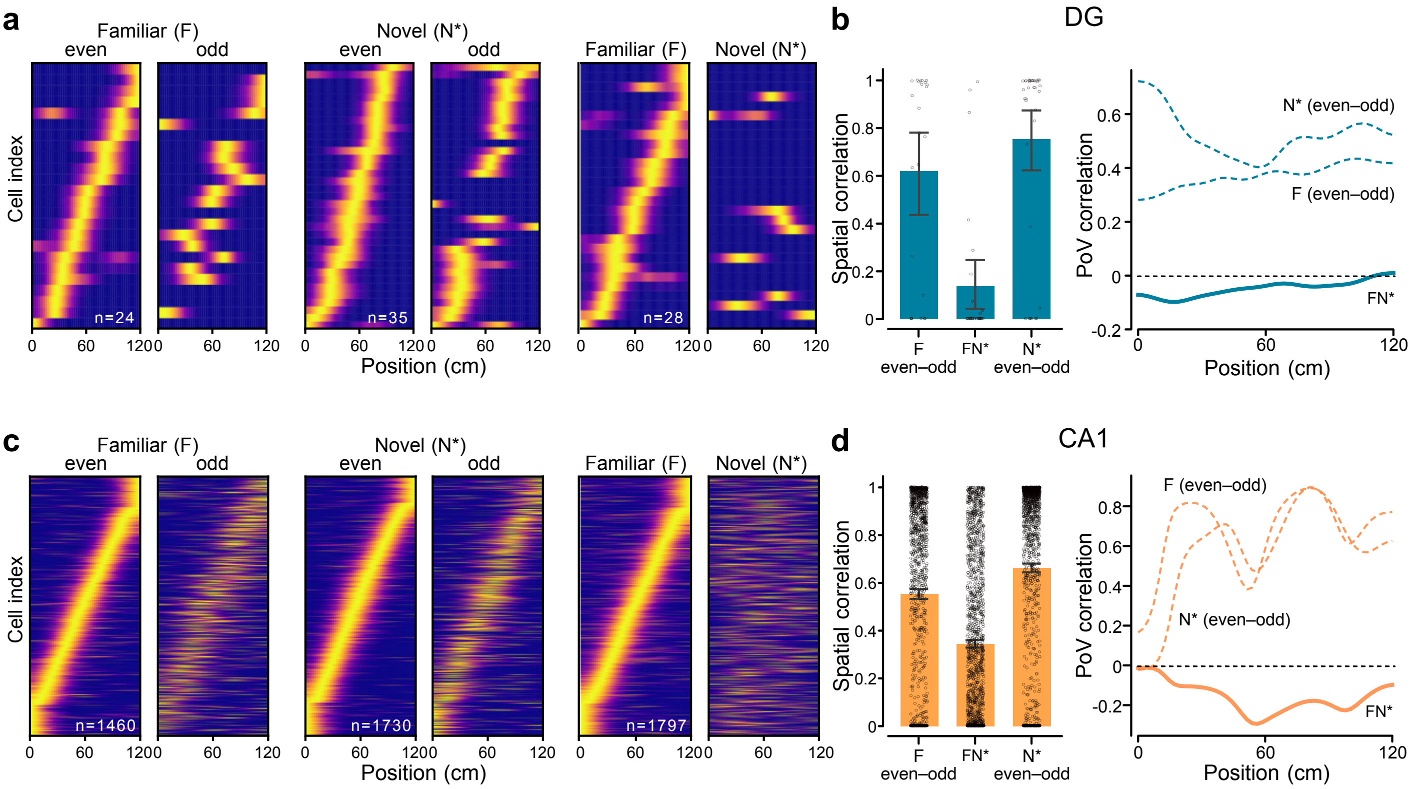


**Extended Data Figure 2**: Spatial activity maps in different environments in the dentate gyrus and CA1

**a,** Spatial activity maps of all spatially modulated cells in familiar (F) or novel (N*) environments sorted by the position of maximal activity (normalized for each cell) in familiar even laps (left), novel even laps (middle) or all familiar laps (right), recorded from the dentate gyrus (5 sessions from 2 mice). **b,** Left: Correlations between mean spatial activity maps across place cells in the same familiar environment (F even-odd, left bar), in different environments (FN*, middle bar), and in the same novel environment (N* even-odd, right bar) in the dentate gyrus (F even–odd: 0.62 ± 0.08, n=24 cells; FN*: 0.13 ± 0.06, n=28 cells; N* even-odd: 0.75 ± 0.07, n=35 cells; 5 sessions recorded from 2 mice). Right: Population vector correlation (PoV) across mean spatial activity maps for all spatially modulated cells against the position of the animals in the virtual corridor. **c,** Same as in **a**, but for CA1 (20 sessions from 7 mice). **d,** Same analysis as in **b**, but for CA1 (left; F even–odd: 0.55 ± 0.01, n=1460 cells; FN*: 0.34 ± 0.01, n=1797 cells; N* even-odd: 0.66 ± 0.01, n=1730 cells; 20 sessions recorded from 7 mice).

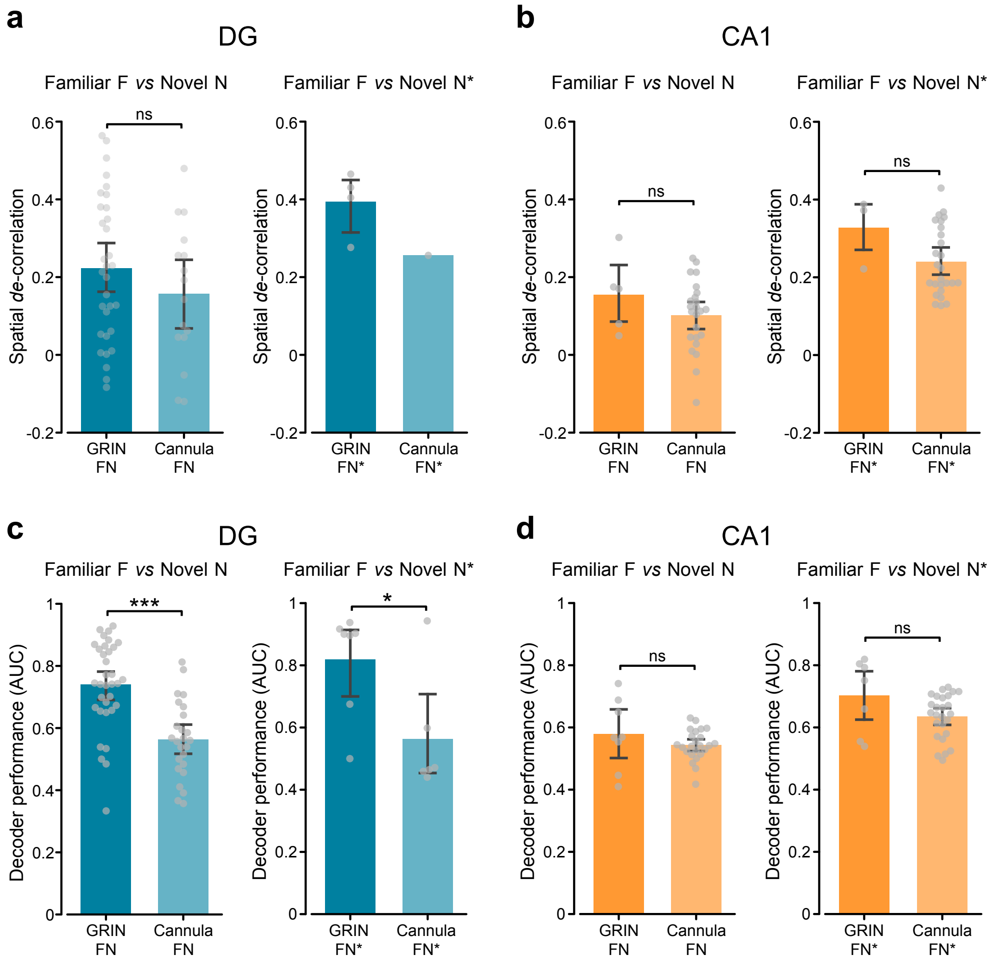


**Extended Data Figure 3**: Comparison of imaging data obtained from animals implanted with GRIN lenses or cannula windows

**a,** Left: Spatial *de*correlation in the dentate gyrus (left, FN; GRIN: 0.33 ± 0.04, n=32 sessions; Cannula: 0.24 ± 0.05; n=16 sessions; P>0.05; right, FN*, GRIN: 0.54 ± 0.05, n=4 sessions; Cannula: 0.37 ± 0.00, n=1 sessions; P>0.05). **b,** Same as **a**, but for CA1 recordings (left, FN; GRIN: 0.15 ± 0.04, n=5 sessions; Cannula: 0.10 ± 0.02, n=24 sessions; P>0.05; right, FN*; GRIN: 0.33 ± 0.02, n=3 sessions; Cannula: 0.24 ± 0.02, n=26 sessions; P>0.05). **c,** Quantification of the decoder performance (AUC) in the dentate gyrus comparing the prediction between environments (left, FN; GRIN: 0.74 ± 0.02, n=34 sessions; Cannula: 0.56 ± 0.02, n=25 sessions; $P<10^{-5}$; right, FN*; GRIN: 0.82 ± 0.06, n=7 sessions; Cannula: 0.56 ± 0.07, n=6 sessions; P<0.05). **d,** Same as **c**, but for CA1 recordings (left, FN; GRIN: 0.58 ± 0.04, n=8 sessions; Cannula: 0.54 ± 0.01, n=24 sessions; P>0.05; right, FN*; GRIN: 0.70 ± 0.04, n=7 sessions; Cannula: 0.64 ± 0.01, n=26 sessions; P>0.05). ns, not statistically significant. * P<0.05; *** $P<10^{-5}$.

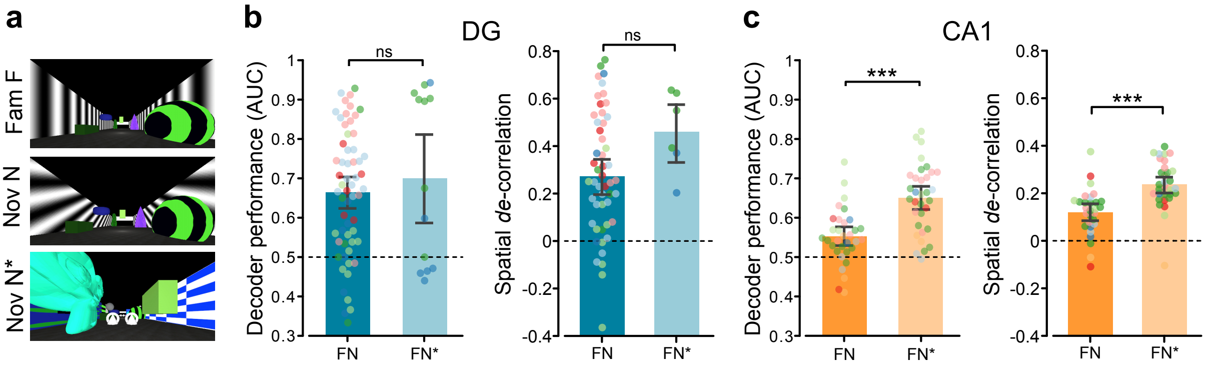


**Extended Data Figure 4**: Neuronal discrimination of small and large changes in the environment

**a,** Views of the virtual environments (Fam (F), familiar; Nov (N) or (N*), novel). **b,** Left: Quantification of the decoder performance in the dentate gyrus comparing the prediction between environments FN and FN*. Dots represent single recorded sessions from different animals (FN, 0.67 ± 0.02, n=59 sessions from 6 mice; FN*, 0.70 ± 0.06, n=13 sessions from 2 mice; P>0.05). Right: Spatial decorrelation, quantified as the difference between spatial correlations within same and different environments (FN and FN*) in the dentate gyrus. Dots indicate single recorded sessions from different animals (FN, 0.30 ± 0.03, n=48 sessions from 6 mice; FN*, 0.51 ± 0.05, n=6 sessions from 2 mice; P>0.05). **c,** Same as **b**, but for CA1 recordings. Left: Dots represent single recorded sessions from different animals (FN, 0.55 ± 0.01, n=32 sessions from 7 mice; FN*, 0.65 ± 0.02, n=33 sessions from 7 mice; $P<10^{-5}$). Right: Dots indicate single recorded sessions from different animals (FN, 0.55 ± 0.01, n=29 sessions from 7 mice; FN*, 0.65 ± 0.01, n=29 sessions from 7 mice; $P<10^{-5}$). ns, not statistically significant; *** $P<10^{-5}$.

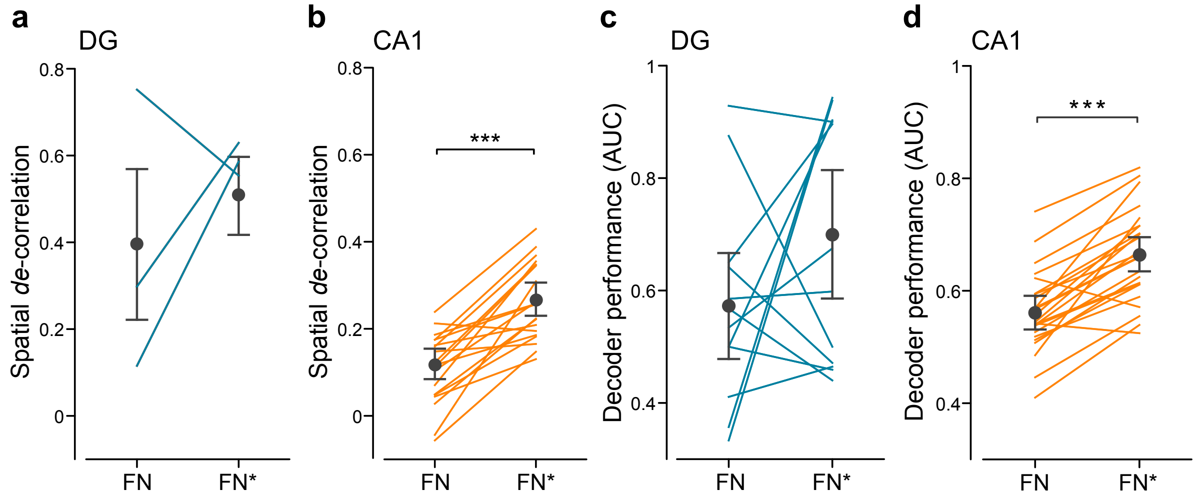


**Extended Data Figure 5**: Paired comparison of neuronal discrimination of small and large changes in the environment

Only paired data obtained in all 3 environments (familiar F, novel N, novel N*) were used in this figure.

**a,** Spatial *de*correlation, quantified as the difference between spatial correlations within the same and different environments (FN and FN*) in the dentate gyrus. Lines indicate paired sessions (FN, 0.39 ± 0.10; FN*, 0.51 ± 0.05; n=3 sessions; P>0.05). **b,** Same as **a**, but for CA1 recordings (FN, 0.12 ± 0.01; FN*, 0.27 ± 0.02; n=19 sessions; $P<10^{-5}$). **c,** Quantification of the decoder performance (AUC) in the dentate gyrus comparing the prediction between environments (FN and FN*). Lines represent paired sessions (FN, 0.57 ± 0.05; FN*, 0.70 ± 0.06; n=12 sessions; P>0.05). **d,** Same as **c**, but for CA1 recordings (FN, 0.56 ± 0.01; FN*, 0.66 ± 0.02; n=24 sessions; $P<10^{-7}$). *** $P<10^{-5}$.

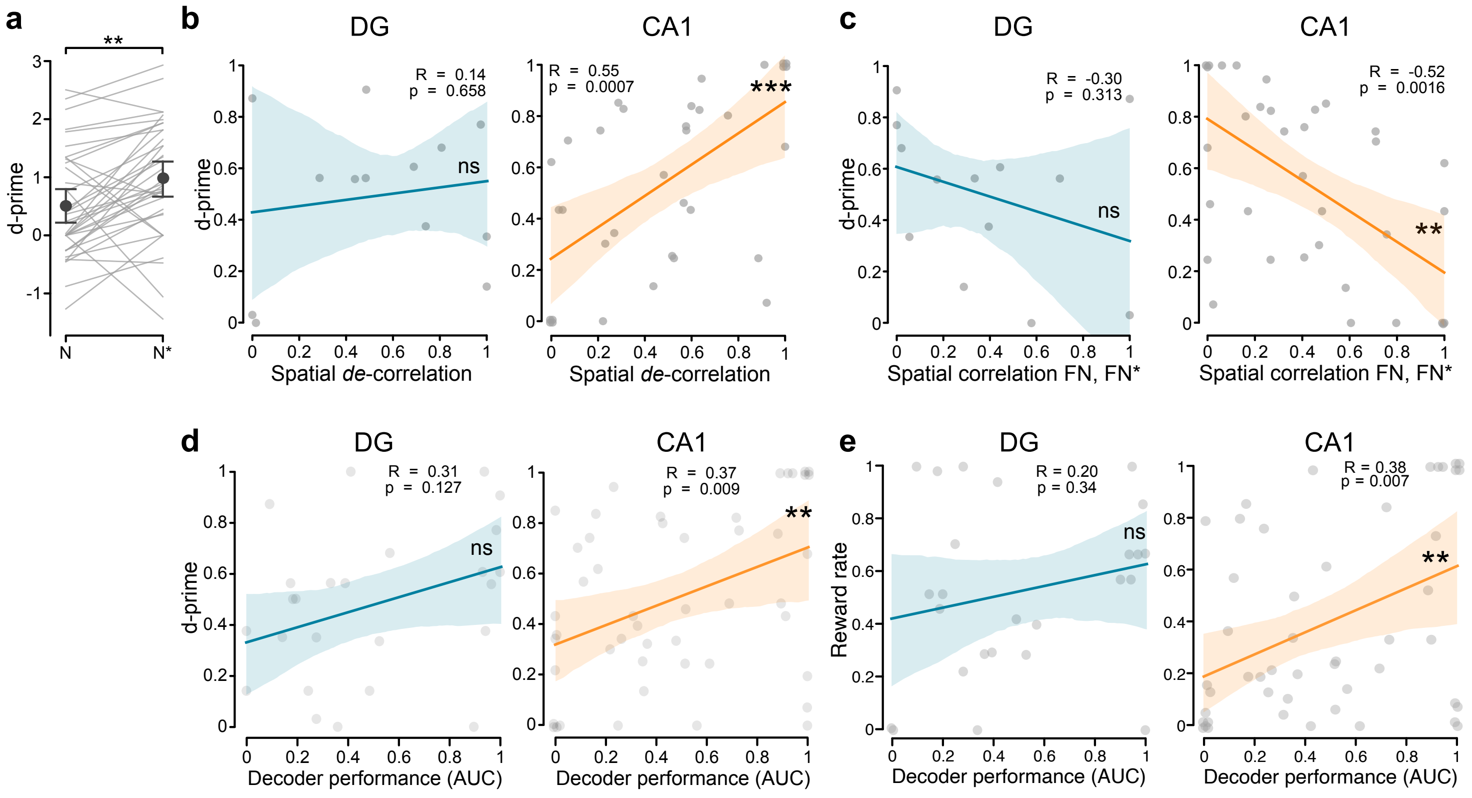


**Extended Data Figure 6**: Neuronal discrimination in CA1, but not in the dentate gyrus, reflects behavioral discrimination

**a,** Supplement to Figure 4c: Quantification of the behavioral performance in the novel (N and N*) environments, measured as d-prime (see Methods). Light grey lines refer to single sessions of all animals, and dark grey dots represent mean ± SEM (N, 0.50 ± 0.15; N*, 0.98 ± 0.16; n=37 sessions; P<0.01). **b,** Pearson's correlation between behavioral performance (d-prime) and spatial *de*-correlation in the dentate gyrus (DG, left: 13 sessions from 2 mice; P>0.05) and CA1 (right: 34 sessions from 7 mice; P<0.001). **c,** Pearson's correlation between behavioral performance (d-prime) and spatial correlation between familiar (F) and novel (N or N*) environments in DG (left: 13 sessions from 2 mice; P>0.05) and in CA1 (right: 34 sessions from 7 mice; P<0.01). **d,** Left: Pearson's correlation between behavioral performance (d-prime) and decoder performance (measured as AUC; see Methods). Light grey dots indicate single recorded sessions from dentate gyrus (DG) implanted animals (25 sessions from 2 mice, P>0.05). Right: Pearson's correlation between behavioral and decoder performance. Dots indicate single recorded sessions from CA1 implanted animals (P<0.1; 48 sessions from 7 mice). **e,** Pearson's correlation between behavioral performance, measured as reward rate (hits per lap) within the novel environments (N and N*) and decoder performance. Dots indicate single recorded sessions from DG (left: 25 sessions from 2 mice; P>0.05) and CA1 implanted animals (right: 48 sessions from 7 mice; P<0.01).

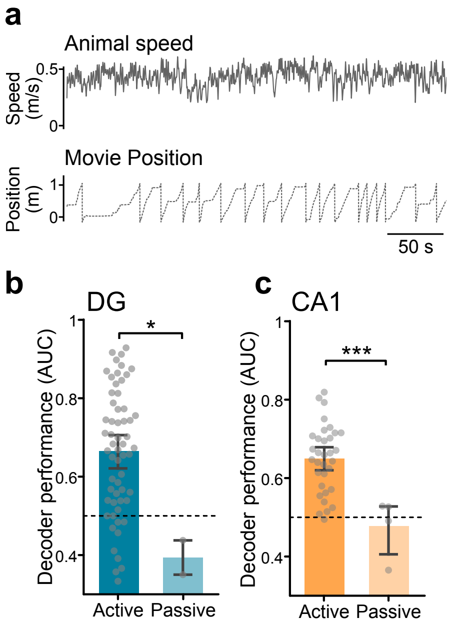


**Extended Data Figure 7**: Hippocampal neuronal discrimination requires active spatial navigation

**a,** Representative open-loop recording session showing the speed of the animal on the running wheel (top trace), and the uncoupled position on the virtual reality track, as imposed by a pre-recorded virtual-reality session (bottom dotted trace). **b,** Quantification of the decoder performance comparing the active and passive spatial navigation conditions. Dots represent single imaging sessions recorded from dentate gyrus (DG) implanted animals (active condition (AUC for FN and FN*): 0.66 ± 0.02, 59 sessions from 6 mice; passive condition (AUC for FN*): 0.34 ± 0.04, 2 sessions from 2 mice; P<0.05). **c**, as in **b**, but for imaging sessions recorded from CA1 implanted animals (active condition (AUC for FN*): 0.65 ± 0.01, 33 sessions from 7 mice; passive condition (AUC for FN*): 0.39 ± 0.04, 4 sessions from 4 mice; P<0.001). *, P<0.05; ***, P<0.001.

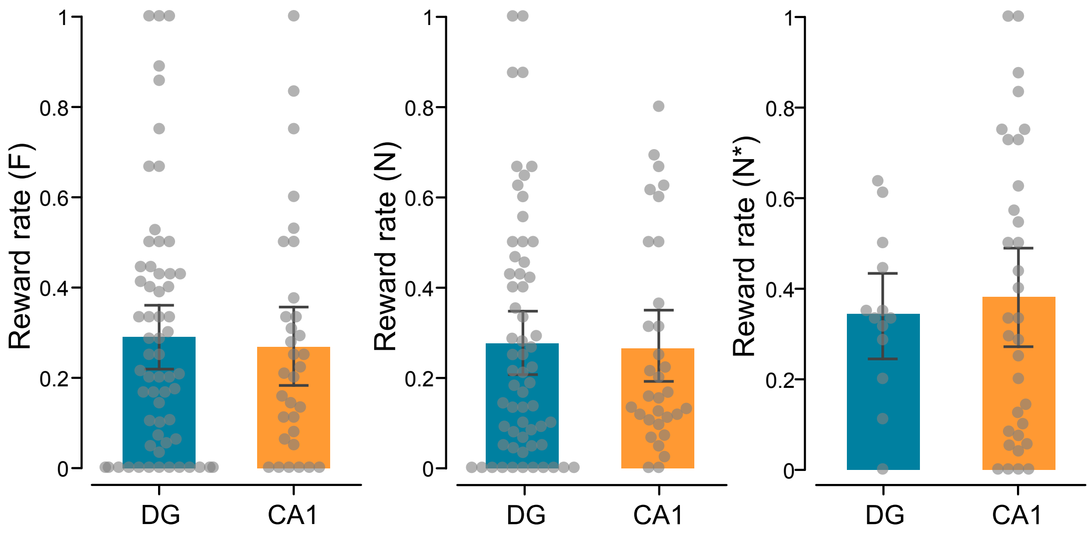


**Extended Data Figure 8**: Similar behavioral performance in animals implanted for CA1 or dentate gyrus imaging

Comparison of behavioral performance (reward rate) in animals implanted for dentate gyrus or CA1 imaging in familiar (F) (left: DG, 0.29 ± 0.04, n=61 sessions from 6 mice; CA1, 0.27 ± 0.05, n=32 sessions from 7 mice; P>0.05), novel (N) (middle: DG, 0.28 ± 0.03, n=61 sessions from 6 mice; CA1, 0.27 ± 0.04, n=32 sessions from 7 mice; P>0.05) and novel (N*) environments (right: DG, 0.34 ± 0.05, n=13 sessions from 2 mice; CA1, 0.38 ± 0.05, n=33 sessions from 7 mice; P>0.05).
